# Investigating mitochondrial gene expression patterns in *Drosophila melanogaster* using network analysis to understand aging mechanisms

**DOI:** 10.1101/2023.05.16.540914

**Authors:** Manuel Mangoni, Francesco Petrizzelli, Niccolò Liorni, Salvatore Daniele Bianco, Tommaso Biagini, Alessandro Napoli, Marta Adinolfi, Pietro Hiram Guzzi, Antonio Novelli, Viviana Caputo, Tommaso Mazza

## Abstract

The process of aging is a complex phenomenon that involves a progressive decline in physiological functions required for survival and fertility. To better understand the mechanisms underlying this process, the scientific community has utilized several tools. Among them, mitochondrial DNA has emerged as a crucial factor in biological aging, given that mitochondrial dysfunction is thought to significantly contribute to this phenomenon. Additionally, *Drosophila melanogaster* has proven to be a valuable model organism for studying aging due to its low cost, capacity to generate large populations, and ease of genetic manipulation and tissue dissection. Moreover, graph theory has been employed to understand the dynamic changes in gene expression patterns associated with aging and to investigate the interactions between aging and aging-related diseases. In this study, we have integrated these approaches to examine the patterns of gene co-expression in *Drosophila melanogaster* at various stages of development. By applying graph-theory techniques, we have identified modules of co-expressing genes, highlighting those that contain a significantly high number of mitochondrial genes. We found important mitochondrial genes involved in aging and age-related diseases in *Drosophila melanogaster*, including UQCR-C1, ND-B17.2, ND-20, and Pdhb. Our findings shed light on the role of mitochondrial genes in the aging process and demonstrate the utility of *Drosophila melanogaster* as a model organism and graph theory in aging research.

## 1. Introduction

Aging can be defined as the time-related deterioration of the physiological functions necessary for survival and fertility [1]. While the specific mechanisms underlying aging are still being studied, it has been shown that mitochondrial DNA (mtDNA) plays a crucial role. In fact, mitochondrial dysfunction, including decreased oxidative capacity and increased oxidative damage, is thought to substantially contribute to biological aging [2,3]. This is due to the higher rate of genomic variants and less efficient repair machinery compared to nuclear DNA: mtDNA’s mutation rate is up to 15 times higher than that of nuclear DNA [4].

For this matter, *Drosophila melanogaster* has been crucial in advancing our understanding of this physiological process and has become a widely used model organism that has distinct advantages in aging research [5]. Commonly known as the fruit fly, it can provide many advantages as a model for research [6] because of the several characteristics that enable it to be widely used in many different applications: low cost of rearing and housing, absence of regulatory oversight for their use in experiments, ease of generating large populations, well defined dietary requirements, easily quantified reproductive output, distinct tissues that can be dissected and genetically manipulated, and furthermore, some of their tissues are equivalent to many of those found in mammals [6]. It is worth remarking that *Drosophila* has significantly helped in studying the hallmarks of aging (genomic instability, telomere attrition, epigenetic alterations, loss of proteostasis, deregulated nutrient sensing, mitochondrial dysfunction, cellular senescence, stem cell exhaustion, and altered intercellular communication [7]). Since 1983, in fact, several studies have been conducted on components of the insulin and IGF-1 like signaling (IIS) pathway in order to genetically manipulate the lifespan of worms and flies with some mutations that were found to be effective in extending their lifespan [8],[9]. This eventually led to the identification of polymorphisms in the insulin pathway transcription factor FOXO associated with the length of life in humans [10]. Furthermore, it has been demonstrated that IGF-1 signaling alters mitochondrial function and enhances their fitness. This includes the maintenance of mitochondrial DNA/RNA [11].

In a biological system, molecules and genes cooperate to perform various biological functions. They engage in dynamic interactions, and graphs are a formal representation of this interaction. In mathematics, a graph is a structure made up of a set of *nodes* or *vertices* and a set of *edges* or *arcs* that show pairwise relationships between the nodes and vertices. The mathematical theory that lies behind graphs has its origins in recreational math problems, but it has now become a significant area of mathematical research with applications in various fields, including computational biology. In computational biology, graph theory is used to represent and analyze complex biological systems such as protein-protein interactions, gene regulatory networks, and metabolic pathways. By representing biological systems as graphs, researchers can use graph algorithms and metrics to study the structural and functional properties of these networks, predict new interactions, and identify key network components that may be targeted for drug development. This discipline has led to many important discoveries, such as the identification of disease-causing genes and the development of new drugs, and has allowed “Network medicine” to grow and develop.

Network medicine is an interdisciplinary field that aims to understand the complexity of biological systems by mapping out the interactions between genes, proteins, and other molecular components. The field uses an integrated, network-based, systems biology-driven approach to define the etiology of complex diseases, reclassify complex diseases, and develop new treatments and preventive strategies [12]. Several studies have investigated the links between network medicine and aging. For example, one study found that the aging process is associated with changes in gene expression networks that affect multiple biological pathways, including metabolism, inflammation, and cell signaling [13]. Another study revealed that caloric restriction, a well-known anti-aging strategy, is connected to decreased DNA damage response (DDR) signaling and senescence burden, possibly via the modulation of gene networks [14]. Moreover, recent experimental evidence has suggested that the genetic or pharmacological ablation of senescent cells, which are known to promote aging and age-related diseases, can extend lifespan and improve health. This approach, known as ‘senotherapy’, is based on the identification of senescence-associated molecular pathways and the development of drugs that target them. Network medicine has been proposed as a useful tool for identifying new targets for senotherapy, as it allows for the integration of multiple sources of data and the identification of the key regulators of senescence networks [13]. Hence, the computational analysis of networks can help identify genes correlated to certain biological pathways, diseases, or phenotypes, which can provide insights into the underlying molecular mechanisms of these processes.

Several computational methods have been borrowed from graph theory over the years to study the dynamics of biological systems [15]. Most often, networks were drawn to represent transcriptional networks, where nodes represented genes and edges represented the strength of co-expression between the genes. These networks were studied to identify “key player” nodes, namely nodes with relevant topological characteristics [16,17], molecular crosstalks [18,19], or functional modules or clusters of genes that co-express under similar conditions [20,21]. These modules, in turn, were often annotated with biological functions, allowing for the identification of pathways and biological processes that were significantly associated with the co-expressed genes.

Various network approaches have been developed over the years, each using different methodologies to detect modules of co-expressed genes [22]. There is evidence, in fact, that cluster analysis of biological networks can help identify key molecular players and pathways involved in the aging process. For instance, a recent study used cluster network analysis to identify subgroups of individuals based on their health and disease phenotypes. The study analyzed data from a large population-based cohort of elderly individuals and found that the identified subgroups had different mortality risks and patterns of health and disease. The authors suggested that this approach can be useful in identifying individuals who are at high risk for certain diseases and in developing personalized interventions [23].

In this study, we combined network, cluster, and gene set enrichment analysis techniques (Supplementary Material F1) to investigate the patterns of gene co-expression in *Drosophila melanogaster* at various developmental stages. We identified modules of co-expressing genes and highlighted those containing significantly high numbers of mitochondrial genes. Therefore, our goal was to use a network-based approach applied to RNA-seq data to find significant mitochondrial genes associated with the aging process in *Drosophila melanogaster*.

## 2. Materials and Methods

### 2.1. Data Sources

*Drosophila melanogaster* data analyzed in this study were retrieved from ENCODE [24] as FASTQ files. Specifically, RNA-Seq data of the whole fly organism were obtained for the following *Drosophila* life stages: embryo (179 samples), larva (69), pupa (64), and adult, distinguished in males (44) and females (41). The reference genome used was *dm6* [25], while the gene annotation file (17.559 genes) was retrieved from FlyBase [26] (GTF: dmel-all-r6.49). Mitochondrial genes for *Drosophila* were retrieved from Mitodrome [27].

### 2.2. Data Preprocessing and Mapping

The quality-control of the raw FASTQ files was performed using the FASTQC software package [28] (v. 0.11.9). Next, sequences were aligned to the *dm6* reference genome using STAR [24,29] (v. 2.7.10a), which was run with default parameters. The resulting read count matrix was filtered by removing genes that were not expressed in any sample; the raw gene counts were normalized into Reads Per Kilobase per Million mapped reads (RPKM) values using the edgeR R package [30]. Normalized data were subjected to Principal Component Analysis (PCA) and the t-distributed Stochastic Neighbor Embedding (t-SNE) method to verify the correct clustering of samples into their respective groups based on their gene expression profiles. The number of principal components was chosen in order to explain 90% of the variance of the gene expression profiles of all samples.

### 2.3. Clustering

Correlation patterns among genes were obtained using the *weighted correlation network analysis* (WGCNA) [31] R software package. Since WGCNA assumes that a gene network obeys a scale-free distribution, it powers the correlation of the genes to an array of soft thresholding values. In this work, we have chosen the power value that, among all, produced the highest similarity with a scale-free network. Therefore, WGCNA yielded an adjacency matrix and a TOM (Topological Overlap Matrix) similarity matrix, the latter being used to perform the actual clustering of samples. Life stages were used as covariates for each sample with the aim of quantifying the module–trait relationships, i.e., the correlations between modules and life stages. This was accomplished through the Pearson correlation coefficient. In order to identify the most representative module of each life stage, we calculated the correlation values between the Module Eigengenes (MEs), defined as the first principal component of a given module, and each life stage.

### 2.4. Node importance

The adjacency matrix resulting from the WGCNA analysis was preprocessed to contain only the top 10% of the scores extracted from the TOM. Then, it was loaded using the *igraph* [27] Python package to make a weighted graph of co-expressed genes. We checked the connectivity for the genes in the module, i.e., the sum of the weights of all the edges connected to a given node. A higher total connectivity implies stronger and more frequent interactions between the nodes and the presence of multiple pathways for information to propagate through the network. Conversely, a lower total connectivity suggests that the nodes are less connected, which may result in fewer interactions and limited avenues for information to flow.

Local topological metrics were computed for nodes in each network module using Pyntacle [32]: (i) *degree* centrality, i.e., the number of edges incident to a node. One way to understand degree centrality is to consider that every edge in a network can be thought of as a path of length 1. Therefore, degree centrality can be seen as a measure that counts the number of these paths that originate from a particular node; (ii) *betweenness* centrality, i.e., the number of shortest paths between all pairs of nodes in the graph that pass through that node. In other words, it measures how often a node appears on the shortest path between any two other nodes in the graph. Nodes with high betweenness centrality are important because they act as “bridges” between different parts of the graph, allowing for efficient communication and the flow of information; (iii) *closeness* centrality, i.e., the average shortest distance from each node to each other node. This is an inverse measure because larger values indicate lower or reduced centrality, as opposed to higher values indicating greater centrality; and (iv) *clustering coefficient*, i.e., the extent to which the neighbors of a node are close to being a clique. If a node has a high clustering coefficient, it suggests that its neighboring nodes are closely connected to each other. On the other hand, if the clustering coefficient of a node is low, it indicates that its neighboring nodes are less connected to each other. The overall clustering coefficient of a network is calculated by taking the average of the clustering coefficients of all the nodes in the network. We also measured the density of the modules by resorting to the *completeness* index [33,34].

A key-player-based analysis was also performed by computing the (i) “Key Player Problem/Negative” (KPP-Neg) index, which gives insights on the contribution of a given node to the cohesiveness of a network (breaking down a network into smaller components, or in cases where fragmentation is not possible, increasing the path lengths between nodes to a degree that they become almost disconnected), and the (ii) “Key Player Problem/Positive” (KPP-Pos) index, which instead focuses on the level of connectivity/embedding of nodes within a network [16], in other words, identifying nodes that can establish direct links or short paths to the maximum number of nodes. Therefore, we computed a KPP-Neg-based measure of network fragmentation, the ^*D*^*F* index, in order to check if the removal of key players leads to a significant increase in the number of disconnected components, and two KPP-Pos-based indices: ^*D*^*R*, which is the weighted proportion of all nodes reached by a set of nodes, which is also known as the “kp-set”, and *m*-reach, which is the count of the number of unique nodes reached by any member of the kp-set in *m* links or less. In this study, we considered only “fast paths” by setting *m* = 3. The key-player analysis was performed on groups of nodes of size 1, namely on individual nodes.

Finally, genes composing the module of interest were subjected to gene set enrichment analysis (GSEA) using PANTHER [35] against the Gene Ontology (GO), which was run with default parameters. To shrink the number of the resulting statistically significant GO terms, a semantic similarity-based clustering approach was used through the REVIGO tool [36]. REVIGO provided a way to summarize the enriched GO terms into more general categories based on their semantic similarities. This strategy yielded a more concise and interpretable summary of the enriched GO terms and helped to identify the biological processes, molecular functions and cellular components that were associated with the genes in the module. A similar analysis was performed with KEGG [37] that regarded the metabolic pathways.

## 3. Results and Discussion

### 3.1. Data Quality and Count Matrix

During the analysis of the RNA sequencing data, the per-sample mean number of reads was found to be 32.051.322, with a wide range of 4.004 to 98.797.584 reads per sample. The average read length was determined to be 88.5, with a range of 76 to 294. In terms of read mapping, it was observed that, on average, 57.41% of the reads across all samples were successfully mapped to the reference genome, with a minimum of 9.42% and a maximum of 94.51%. After filtering out genes that were not expressed in any of the samples, 4.8% of genes (849 out of 17559) were removed from the resulting read count matrix.

### 3.2. Life Stages and Clustering

According to the results of the PCA analysis, 30 principal components were needed to explain 70% of all the variance that the gene count matrix was able to capture (see Figure 1a). As a result, data projected on these components and subsequently subjected to spectral embedding revealed clear contiguity of the major life stages as well as tight clustering of intra-stage groups (i.e., prepupa, pupa, and three instar larval stages) (Figure 1b).

**Figure 1.**
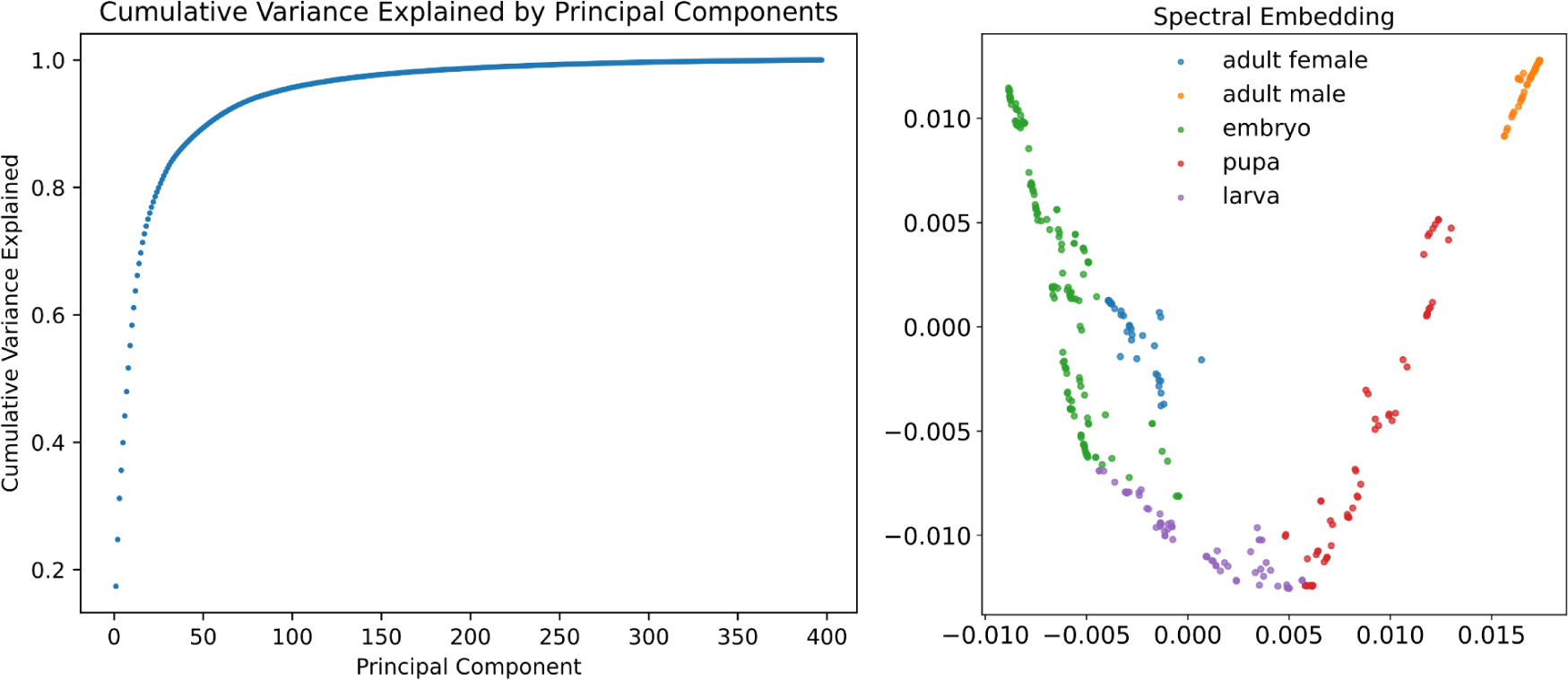
**(a)** Cumulative variance explained as the number of principal components increases; **(b)** Spectral embedding of the expression values of all Drosophila melanogaster samples labeled with the corresponding life stage

Expression data were then processed by WGCNA that was run with default parameters: the power that produced a higher similarity with a scale-free network was 9 (Figure 2a left), and clustering was performed using the resulting TOM by merging modules whose expression profiles were very similar as measured by the correlation of their eigengenes (Figure 2b).

**Figure 2.**
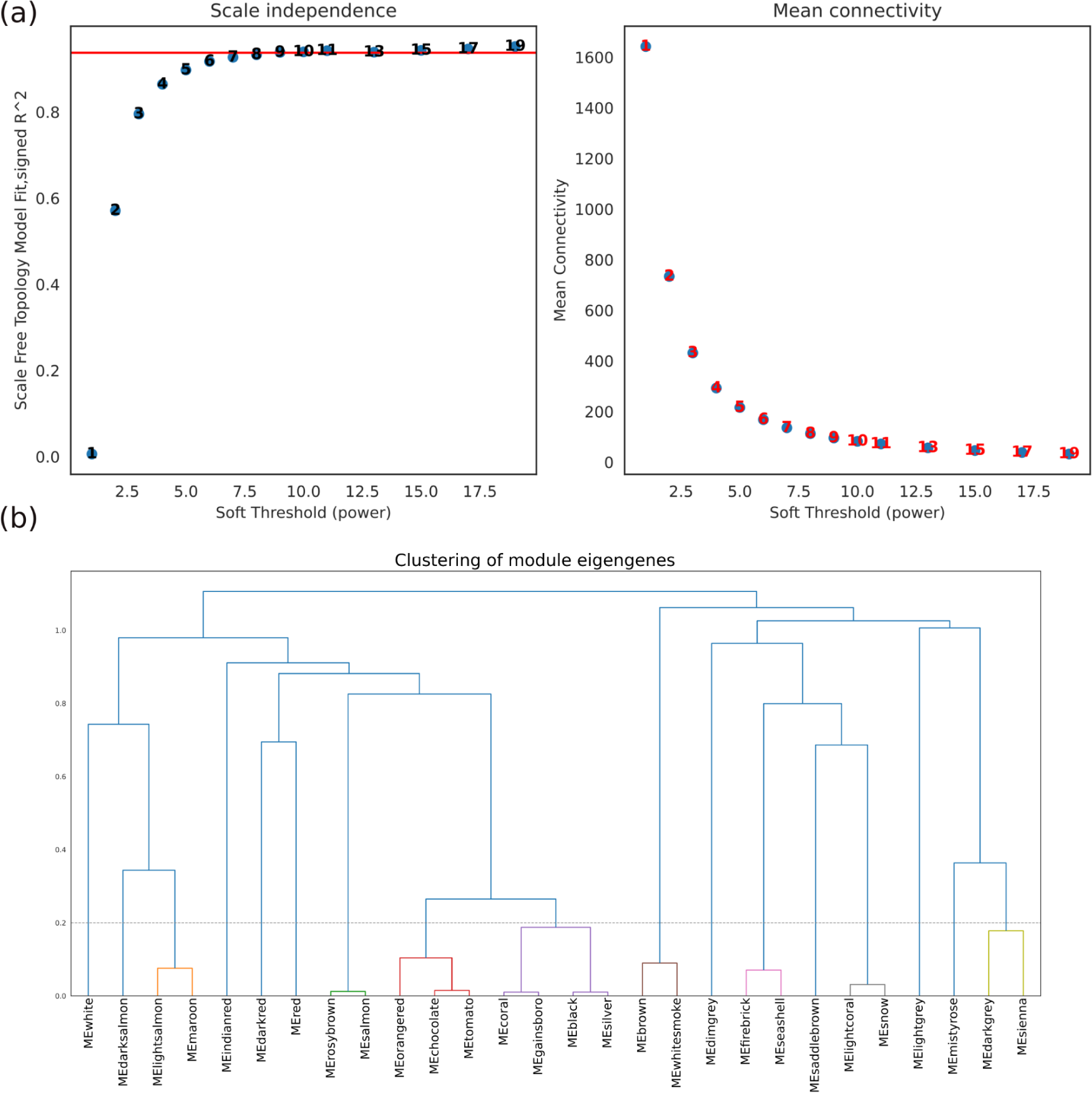
Results of the WGCNA preprocess: **(a, left)** scale-free fit index and mean connectivity (degree, **right**) with respect to the soft-thresholding power (X-axis). The left panel demonstrates the relationship between the scale-free fit index and soft-thresholding power, while the right panel displays the mean connectivity as a function of the soft-thresholding power; **(b)** Clustering dendrogram of genes, with dissimilarity based on topological overlap, together with assigned merged module colors and the original module colors.

Originally, the detected modules were 28; merging the ones whose distance was less than 0.2 gave us the 17 clusters considered in this work.

### 3.3. Module-Trait Relationships

Module-trait relationships were then represented by a heatmap (Figure 3), which summarized the correlation values between the co-expressed genes and each specific life stage. Interestingly, the “black” and “orangered” clusters were strongly positively correlated with the adult male cluster (0.85, *p-value*<0.05; 0.84, *p-value*<0.05 respectively) and negatively correlated with the embryo (−0.54, *p-value*<0.05; -0.38, *p-value*<0.05 respectively). Overall, co-expression levels increased from the embryo to the adult (male) stage in both modules, while they remained almost similar in the females. In this regard, it is interesting to note that genes co-expressed during the embryonic phase are similar to the ones in the adult female, generating similar scores in almost all clusters. The “white” cluster shows a strong positive correlation with the larval cluster (0.52, *p-value*<0.05), but negative correlations in the embryonic and pupa clusters (−0.19, *p-value*<0.05; -0.28, *p-value*<0.05 respectively); a similar behavior is noted for the red cluster, with the exception of the pupa positive correlation (0.58, *p-value*<0.05). Another case regards the “saddlebrown” module, which does not show a drop in correlation from the embryo phase to larval, and it remains low for the males (adult) but has a slight increase for the females (adult).

**Figure 3.**
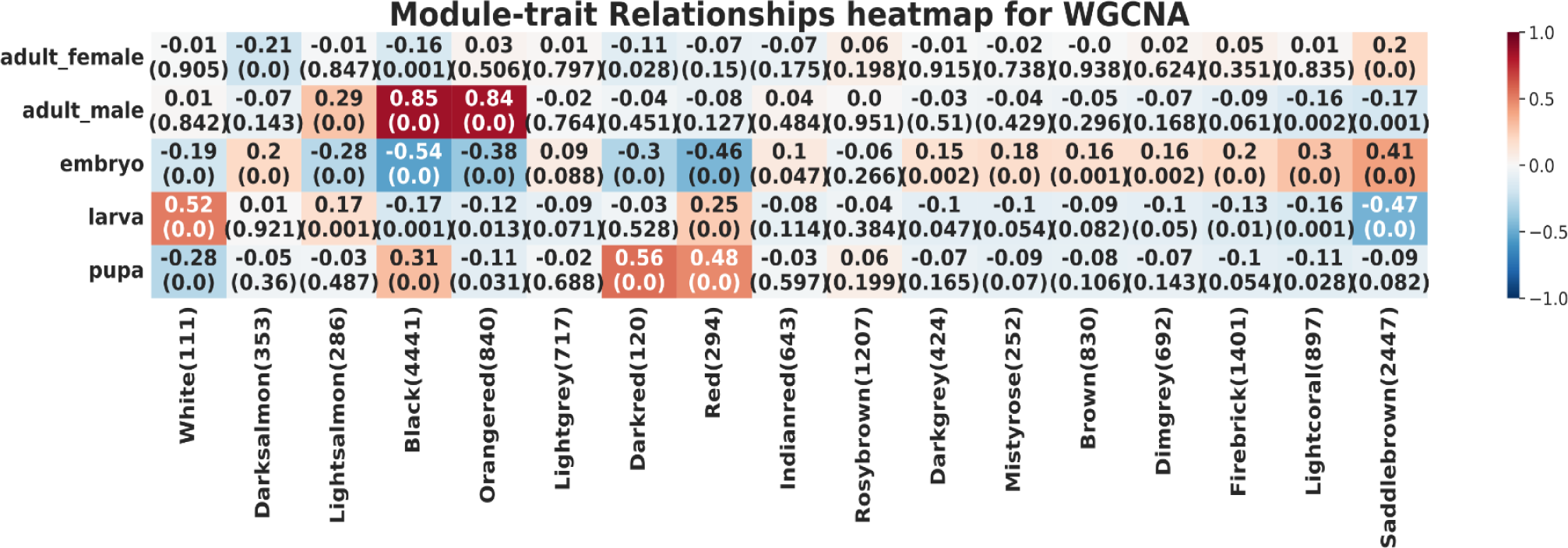
Relationships between consensus modules and life stages. Each column corresponds to a cluster, and each row to a life stage. The numbers in the table report Pearson correlations and p-values in parentheses (meant as the probabilities that one would have found the current results if the correlation coefficients were in fact zero, i.e., null hypothesis), where shades of red show a positive correlation and blue shades a negative correlation.

### 3.4. Network analysis

In order to understand the role of mitochondrial genes in aging, we focused on a cluster that showed a high correlation for any specific life stage and had a higher concentration of mitochondrial genes. As shown in Figure 4, the “lightsalmon” cluster (Supplementary Material F2), despite being a small cluster, contains a high number of mitochondrial genes and shows a positive correlation in the adult male (0.29, *p-value*<0.05), while a negative one in the embryonic phase (−0.28, *p-value*<0.05).

**Figure 4.**
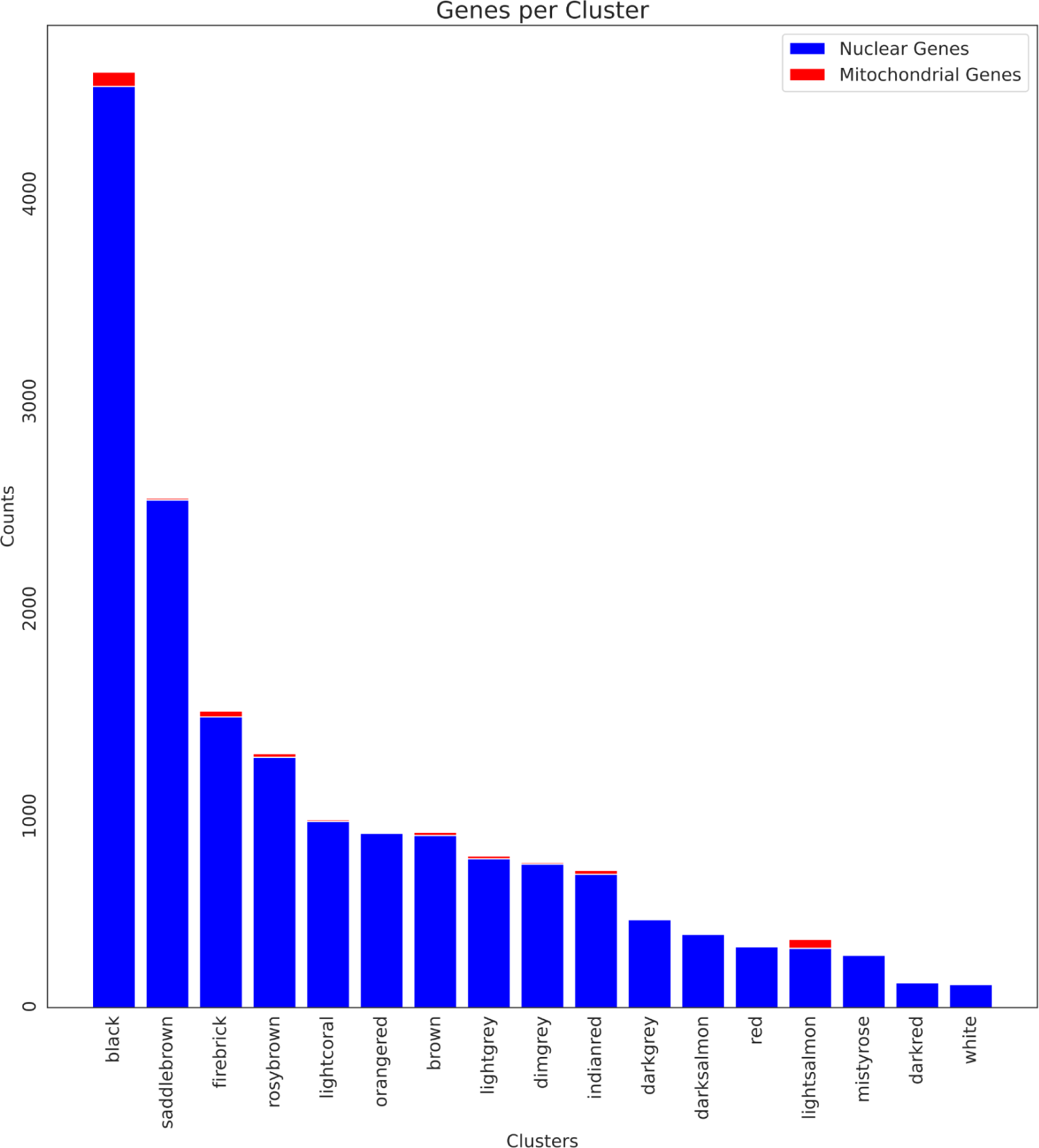
Barplot showing the number of nuclear (blue) and mitochondrial (red) genes for each cluster. The lightsalmon cluster has the highest ratio of mitochondrial genes.

The cluster contains 286 genes, and 43 of them are mitochondrial. We studied its topological organization using Pyntacle. Several local topological metrics (i.e., degree, betweenness, closeness, and clustering coefficient) were calculated starting from the adjacency matrix generated by WGCNA that was cleaned from values belonging to the lowest ninetieth percentile. This filtering step was adopted to minimize noise and density in the resulting network; the 286 nodes of the network were then wired with 3947 edges, which then represented only strong correlations between genes. This network was made up of 28 components.

First, we assessed the connectivity of each node in the cluster: the mean connectivity for the whole network (2.98) was smaller than the average among all the mitochondrial genes, which is 4.46 (Figure 5). This means that mitochondrial genes tend to be more connected to other nodes in the network: among the top 50 genes, 14 of them were mitochondrial genes, and in the top 10, the mitochondrial ones were 4. The two most connected genes are both mitochondria, i.e., CG7430 (13.58) and UQCR-C1 (ubiquinol-cytochrome c reductase core protein 1) (12.76). Other relevant genes are mAcon1 (5th) and Pdhb (Pyruvate dehydrogenase E1 beta subunit) (8th).

**Figure 5.**
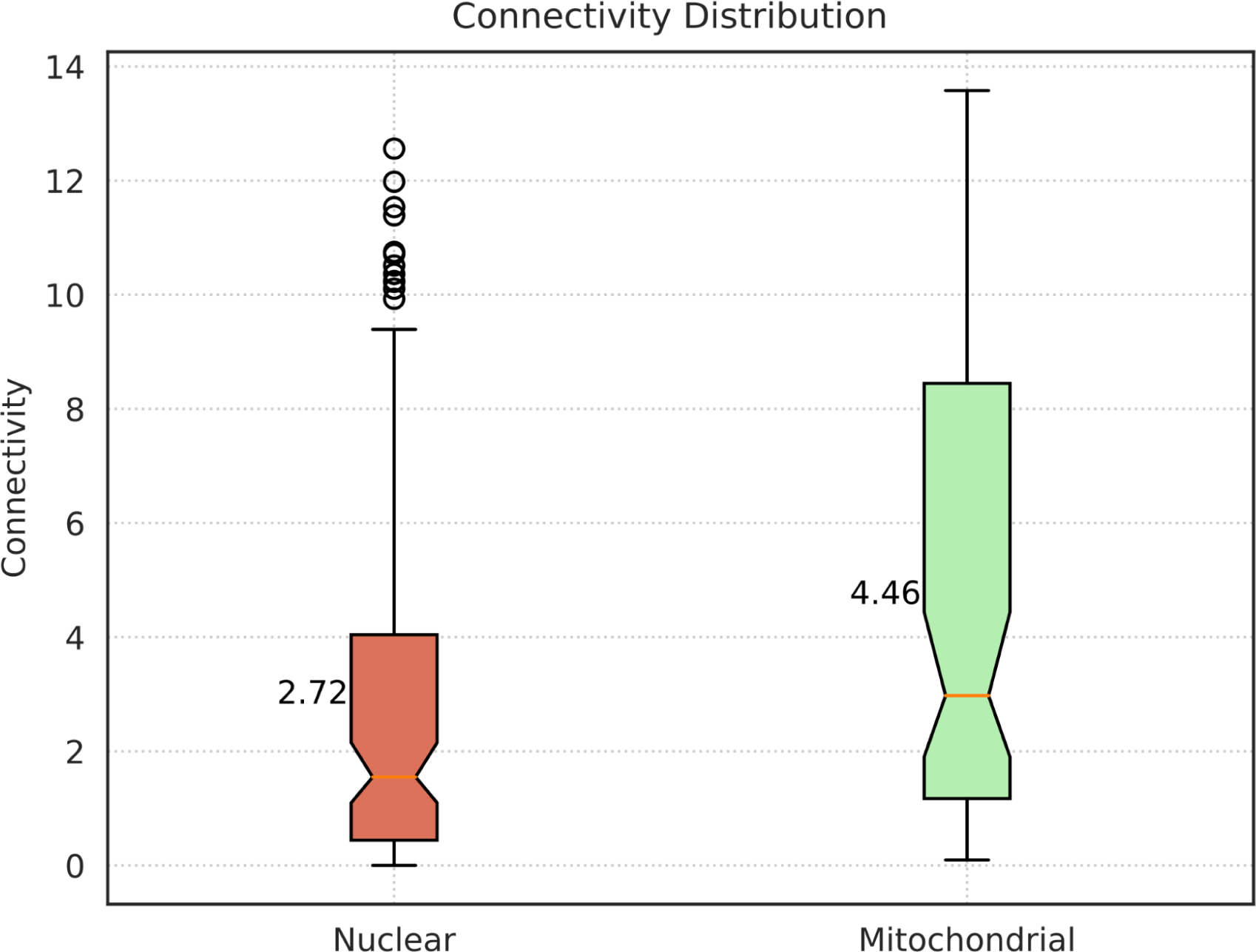
Boxplot showing the distribution of the connectivity in both nuclear and mitochondrial genes for the lightsalmon module. The orange line shows the median, while the reported value is the mean. Mitochondrial genes tend to be more connected.

Then, we evaluated the local centrality measures of genes. As for *degree* centrality, the top 50 genes exhibited an average degree of 67.24, which is higher than the average degree of the whole network (27.60); interestingly, 10 of them were mitochondrial genes. UQCR-C1 was the top mitochondrial gene.

For what concerns the *betweenness* centrality, only 5 mitochondrial genes were included in the top 50; the top mitochondrial gene (mAcon1) among them exhibited a betweenness centrality value of 3738 versus an average for the whole network of 486.12, which means that this gene may play a crucial role in maintaining the connectivity of the network and that, if it were removed, it would significantly disrupt the organization of the network.

Instead, no mitochondrial genes were within the top 50 in terms of closeness, which in a network means that it may take longer for information to flow from that node to other nodes or, in a co-expression network, that they tend to cluster together and are less connected than nuclear genes; however, the top 2 mitochondrial genes as for closeness, i.e., Idh3b and mAcon1, were respectively 0.5 and 0.7 far from the top 50.

As the last centrality measure, we focused on the clustering coefficient; seven mitochondrial genes, i.e., Cchl (Cytochrome c heme lyase), SdhB (Succinate dehydrogenase, subunit B), ND-B17.2 (NADH dehydrogenase B17.2 subunit), CG1907, Hmgcl (3-Hydroxymethyl-3-methylglutaryl-CoA lyase), ND-20 (NADH dehydrogenase 20 kDa subunit), and CG1673, were among the 56 genes with the maximum score, i.e., clustering coefficient = 1. This means that they exhibit a community-like structure with their neighboring genes in the network (mostly other mitochondrial genes). On the contrary, only 3 mitochondrial genes, i.e., aralar1, Hsc70-5, and mEFTu1 (mitochondrial translation elongation factor Tu 1), had a score of 0, meaning that they were loosely connected with their neighborhood.

Finally, we conducted a key-player analysis. It included an assessment of the fragmentation and reachability properties of the network under examination. Interestingly, the network showed high levels of resilience towards fragmentation (quantified by the ^*D*^*F* index), as all nodes gave a score of 0.2 (^*D*^*F* ranges from 0, minimum fragmentation, to 1), which is common in biological networks due to their renowned robustness to exogenous interferences [38–40]. On the other hand, the *m-reach* index showed that mitochondrial genes exhibit high reachability: the mean *m-reach* value for the whole network was 77.04 ∓ 69 std, while, on average, the mitochondrial genes exhibited an average *m-reach* of 127.15 ∓ 57 std. This might be due to the fact that those genes tend to cooperate and cluster together much more than their nuclear counterparts. ABCB7, UQCR-C1, CG7430, and Pdhb were the top mitochondrial genes. Finally, considering the *distance-weighted reach* (^*D*^*R*) index as well, we calculated an average ^*D*^*R* of 0.326 for the mitochondrial cluster (Figure 6), which was almost the same as the whole network (0.3); the whole-network mean also considers isolated nodes, which have ^*D*^*R*=0, by definition. This shows that even if mitochondrial genes tend to form dense communities and cliques, they still maintain a high reachability, on average, to the whole network, which is at least comparable with that of the nuclear genes.

**Figure 6.**
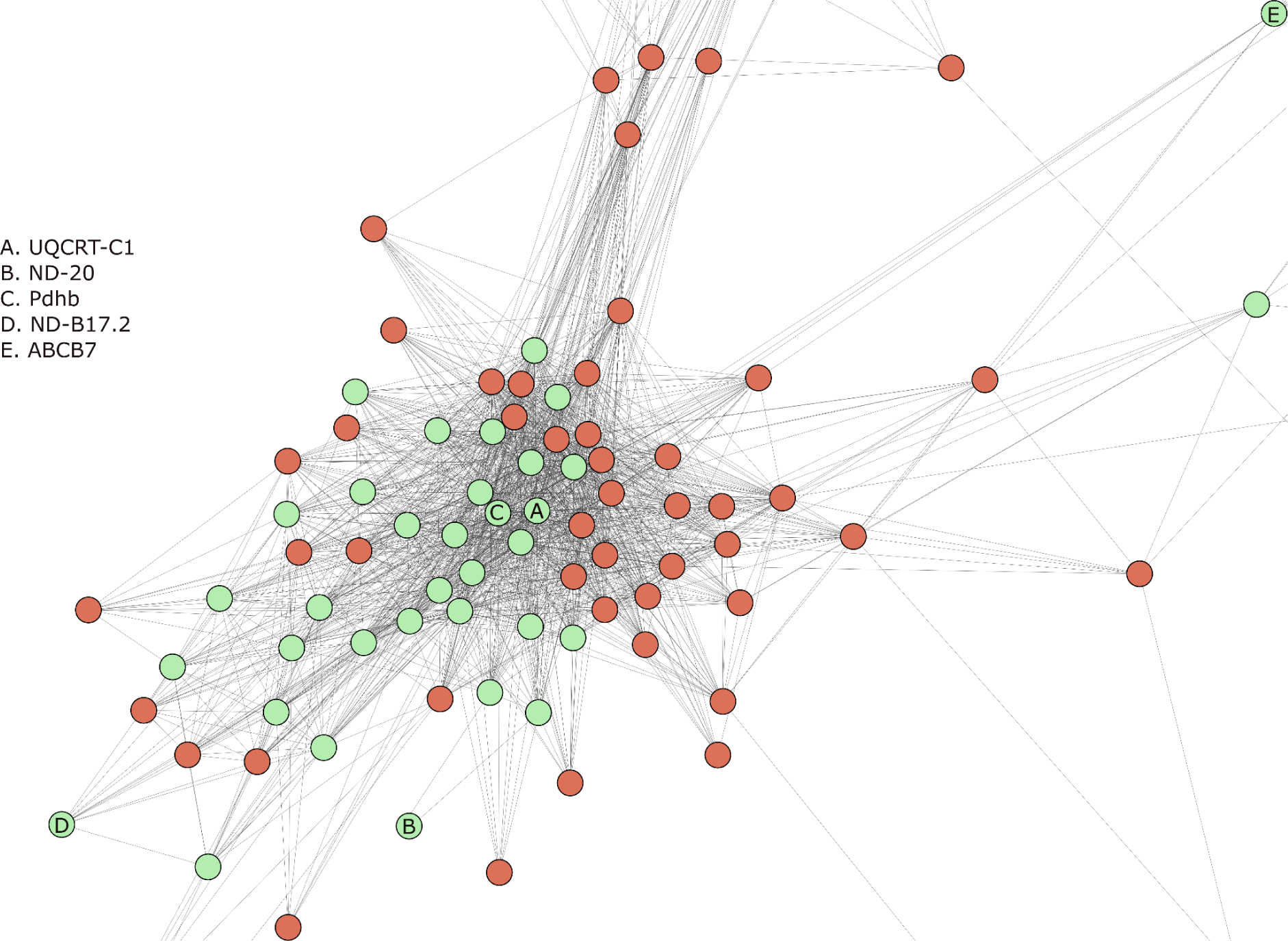
Graphical representation of the “lightsalmon” cluster obtained using *igraph*. Each node is a gene. When two genes co-express, an edge connects them. The “lightsalmon” module contains five topologically highly ranked mitochondrial genes: UQCR-C1, ND-20, Pdhb, ND-B17.2, and ABCB7. Mitochondrial genes are colored in green.

### 3.5. Gene Set Enrichment Analysis

A final step was to study the biological processes, cellular components, and molecular functions in which the 286 genes making up the network module under investigation were involved, with a particular focus on mitochondrial functions. As shown in Figure 7, the genes in this module exploit different functions, the largest part of which regard regulatory and structural processes (light-yellow, ocher). However, five tiles, which are colored “salmon”, significantly enrich mitochondrial-specific functions, thereby confirming the statistical over-representation of mitochondrial genes in this network module. Moreover, since mitochondria are responsible for energy production via oxidative phosphorylation, ion transport across mitochondrial membranes is essential for their proper function and plays a critical role in various cellular processes such as cytoplasmic homeostasis, energy generation, and the compartmentation of metabolism. Several ion transport-related functions were statistically enriched by this analysis and are represented in the figure below by the green tiles. Finally, mitochondria provide the energy required for muscle contraction, as they are responsible for producing ATP. Most ATP is produced in mitochondria through a series of reactions known as the Krebs cycle. During muscle contraction, calcium ions are released, which bind to proteins that enable the myosin and actin filaments to slide past each other, resulting in muscle contraction. In this regard, several muscle and locomotion-related biological processes were statistically enriched and represented by “darkorange” tiles in Figure 7 (Supplementary Material F3). These are particularly important since mitochondrial dysfunction has also been linked to various muscle-related diseases, including mitochondrial myopathies [41].

**Figure 7.**
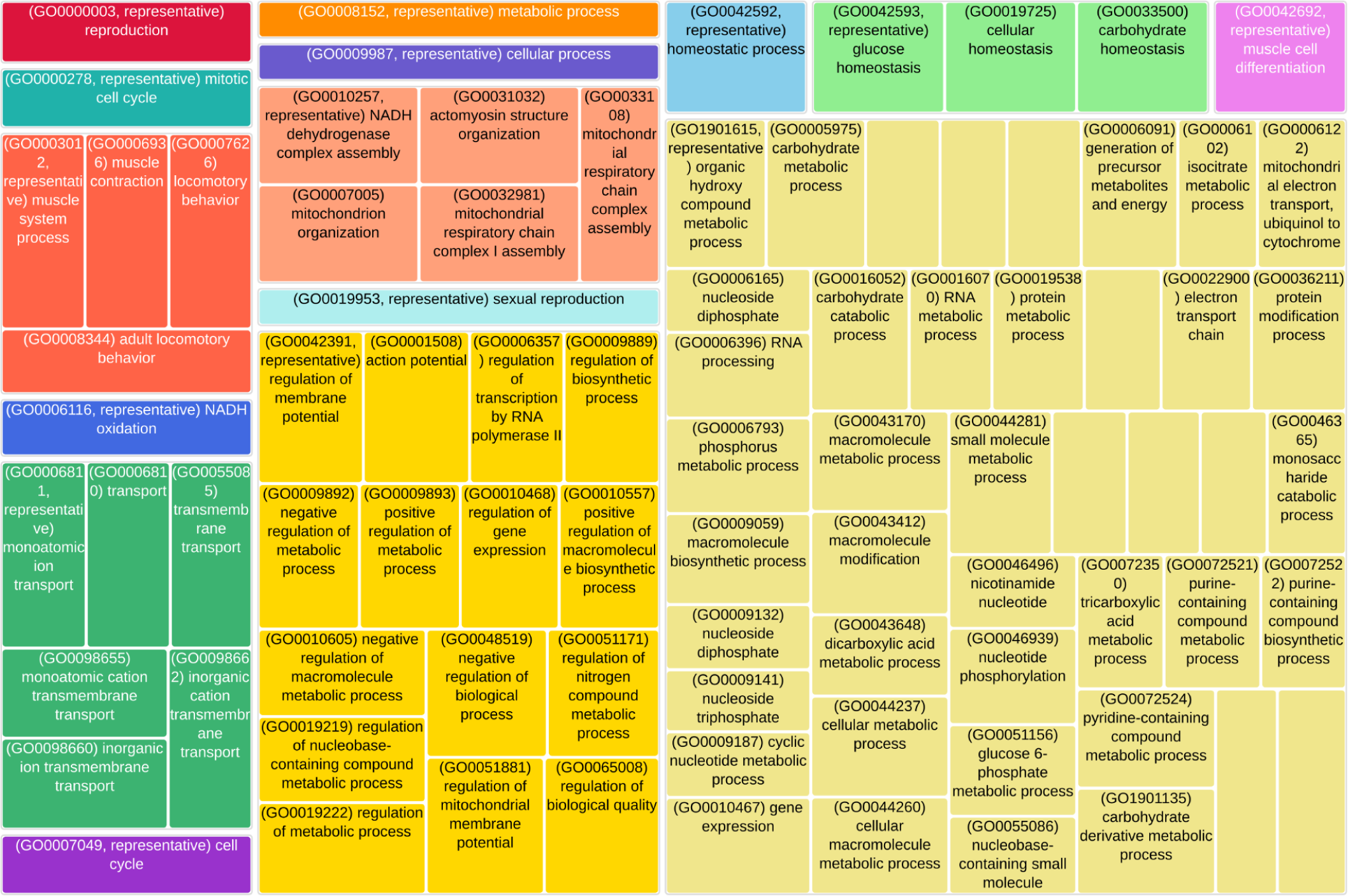
TreeMap representing the GO terms statistically enriched by the genes composing the “lightsalmon” network cluster. The terms, i.e., the TreeMap tiles, are joined into ‘super tiles’ of loosely related GO terms, visualized with different colors. The size of each tile represents either the p-value, or the frequency of the GO term in the underlying GO database.

The analysis of the “Cellular component” GO class of terms confirmed these results (Supplementary Material F4). The most statistically significant GO term was “mitochondrial inner membrane;” which indicates that a significantly high number of genes in our cluster lie in this cellular compartment. Other cellular components resulted from this analysis to be significantly over-represented by the genes of this cluster: the “organelle inner membrane”, “mitochondrial protein-containing complex”, and the “inner mitochondrial membrane protein complex”. These components are essential for maintaining cellular homeostasis, energy production, and metabolic regulation.

The analysis of the GO molecular functions category (Supplementary Material F5) revealed that the genes in this cluster primarily exploit the transport of ions and cations as well as membrane functions associated with ATP synthesis.

In addition, we performed a metabolic pathway enrichment analysis against the KEGG database (Figure 8). Coherently with the previous results, the most significant results relate to mitochondrial processes that were strongly statistically enriched and significant, such as oxidative phosphorylation, carbon metabolism, and the citrate cycle (TCA cycle).

**Figure 8.**
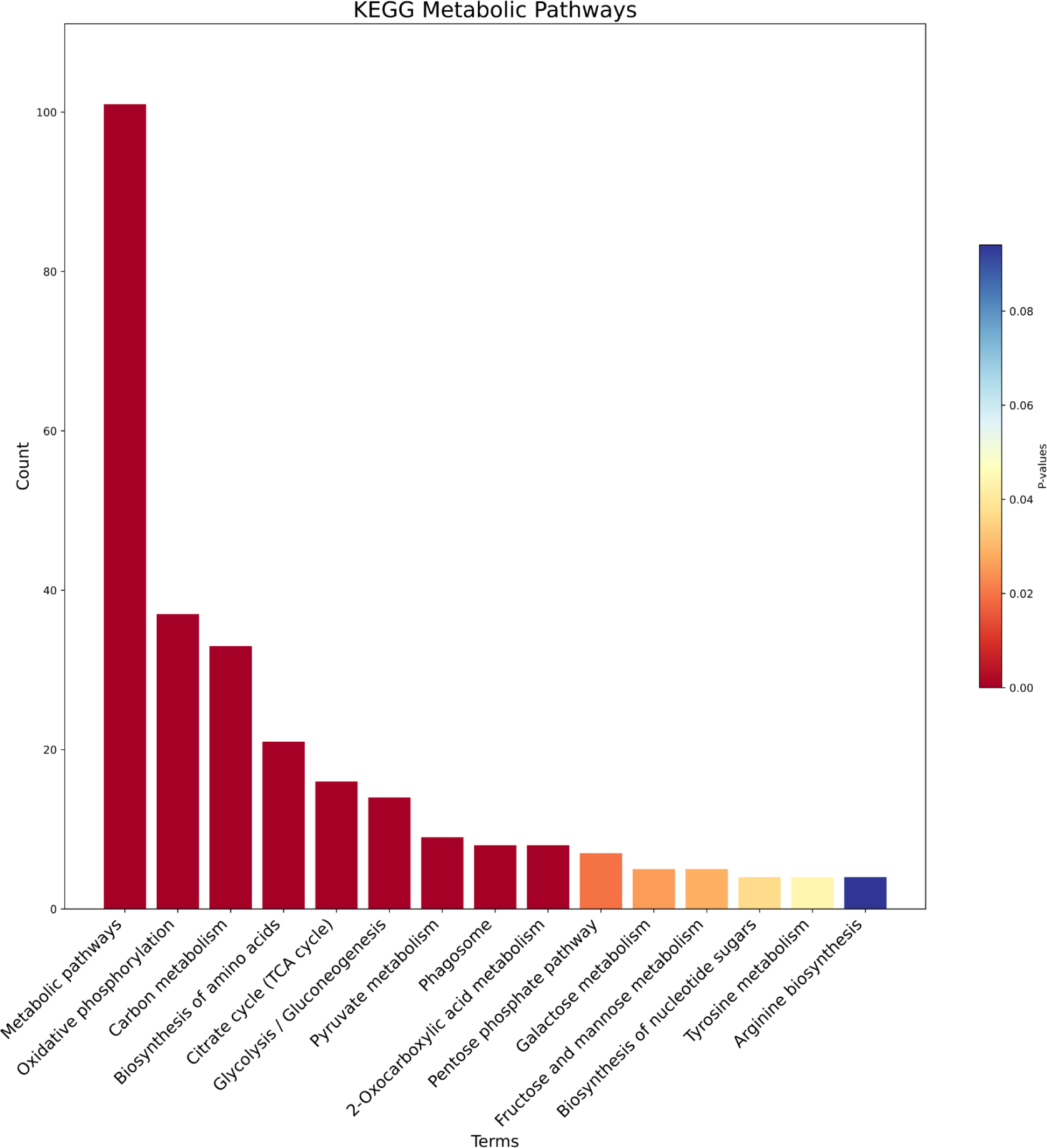
The barplot shows the enriched metabolic pathways from KEGG; for each pathway, we show the gene count (y-axis) and the corresponding p-value (color gradient), where shades of red show a smaller value and blue shades a higher value.

### 4. Concluding remarks

In this study, we showed that network-based methods, particularly when used with high-throughput gene expression data, have the potential to reveal significant insights. In this instance, we used RNA-seq data and a technique called Weighted Gene Co-expression Network Analysis (WGCNA). Using this methodology, previous studies have produced significant findings, such as the identification of potential biomarkers for ischemic stroke. [42][43]. In this work, we have analyzed the gene expression profiles of *Drosophila melanogaster* at different life stages and identified some mitochondrial genes that may have an important role in the process of aging in this model organism: UQCR-C1, which is an important mitochondrial gene in the network from a topological standpoint: it is expressed in several structures, including the adult heart, embryonic/larval midgut, and spermatozoon. It has been investigated in different studies of Parkinson’s disease, and its human ortholog is also involved in Alzheimer’s disease and multiple mitochondrial dysfunction syndrome. ND-B17.2 is known to be involved in the mitochondrial respiratory chain, and its human ortholog has been studied regarding nuclear type mitochondrial complex I deficiency 23 (MC1DN23). ND-20 has been studied as a part of the mitochondrial electron transport chain and, particularly, is involved in the determination of life span. Knockdown of Pdhb, instead, has been associated with a shortened lifespan in adult flies [44].

In *Drosophila*, homozygous variants of mAcon1 are lethal, and knockdown of the protein through RNA interference has been found to cause reduced locomotor activity, a shortened lifespan, and increased cell death in the developing brain. These findings suggest that mAcon1 is essential for proper mitochondrial activity, energy metabolism, and aging [45].

On this path, we aim to further investigate the role of these mitochondrial genes in aging and age-related diseases and plan to expand our network-based approach to other model organisms, such as other *flies and insects*, and datasets to identify conserved molecular mechanisms underlying aging and age-related diseases.

## Supporting information

Supplementary Material F1. Graphical representation of the analysis pipeline used in this work.

Supplementary Material F2. Whole network representing the "lightsalmon" cluster; nuclear genes are in red, while mitochondrial genes are in green.

Supplementary Material F3. Dot plot of the top 10 Biological Processes resulting from the GSEA analysis.

Supplementary Material F4. Dot plot of the top 10 Cellular Components resulting from the GSEA analysis.

Supplementary Material F5. Dot plot of the top 10 Molecular Functions resulting from the GSEA analysis.

## Notes

### Competing Interest Statement

The authors have declared no competing interest.

### Summary of Updates

We fixed some minor issues in he text. Improve quality of the figures and expanded captions. We expanded the result and discussion section.

