## Supplementary figures and images for "Investigating mitochondrial gene expression patterns in *Drosophila melanogaster* using network analysis to understand aging mechanisms"

### Supplementary Material F1.png

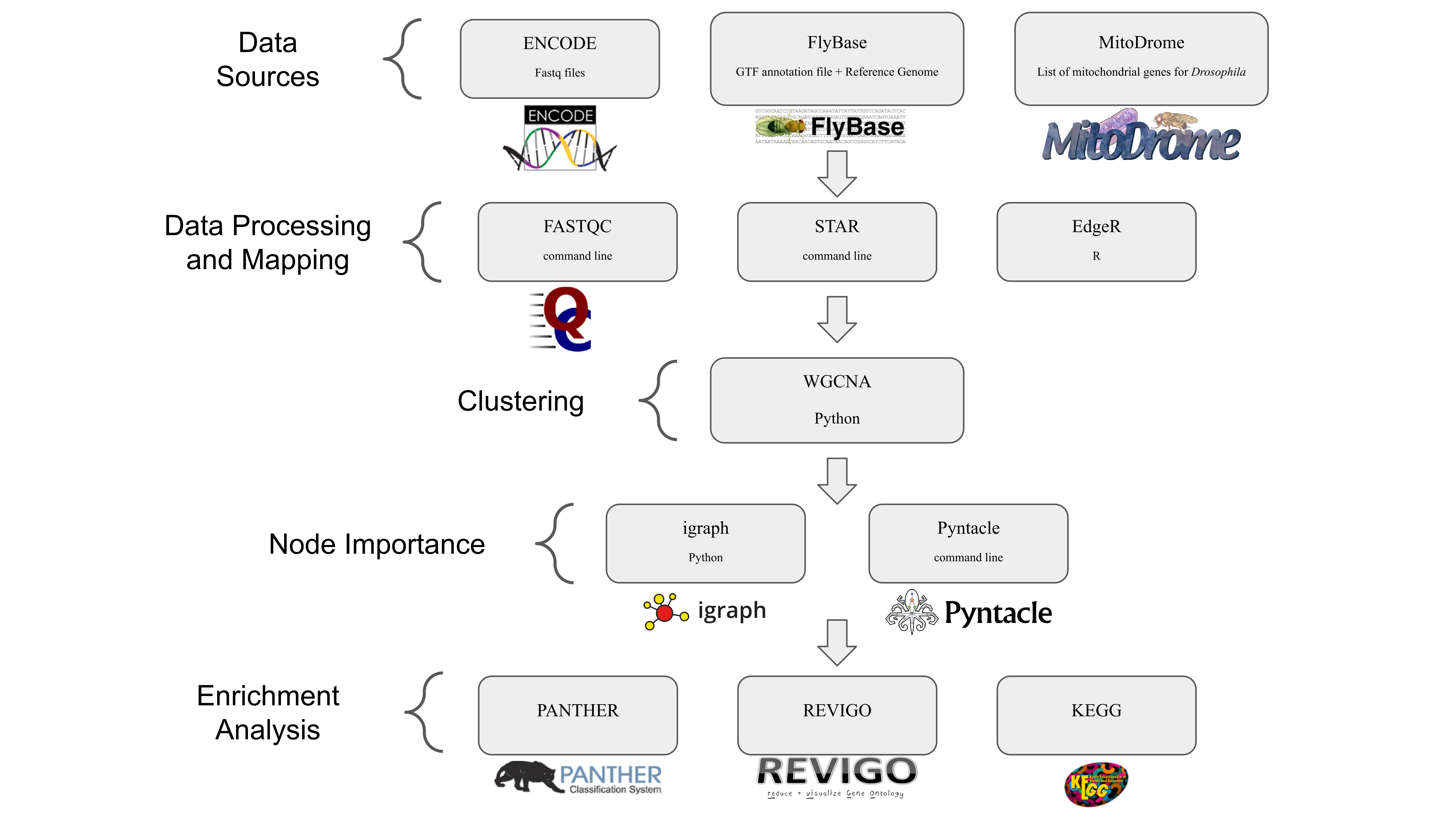

### Supplementary Material F2.png

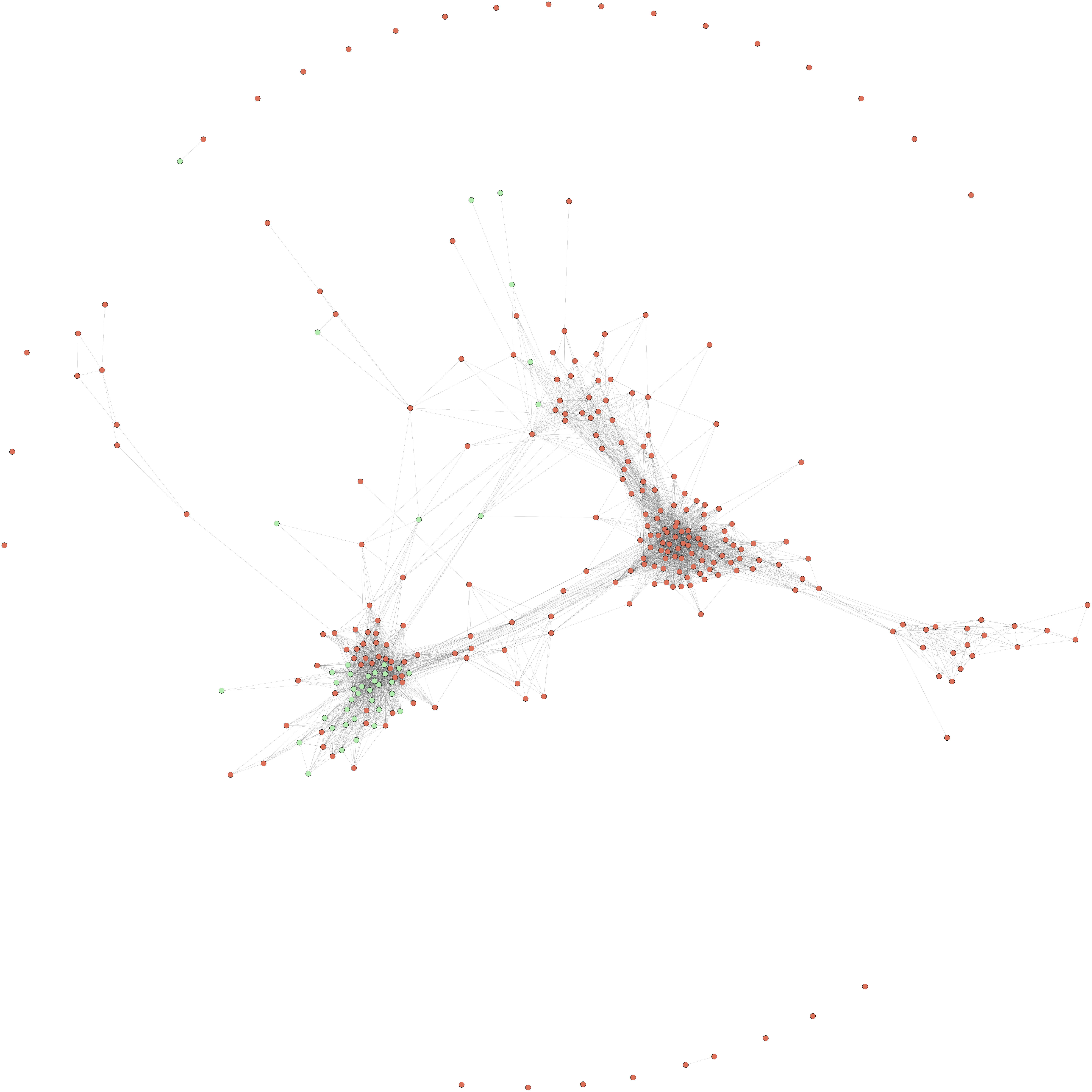

### Supplementary Material F3.png

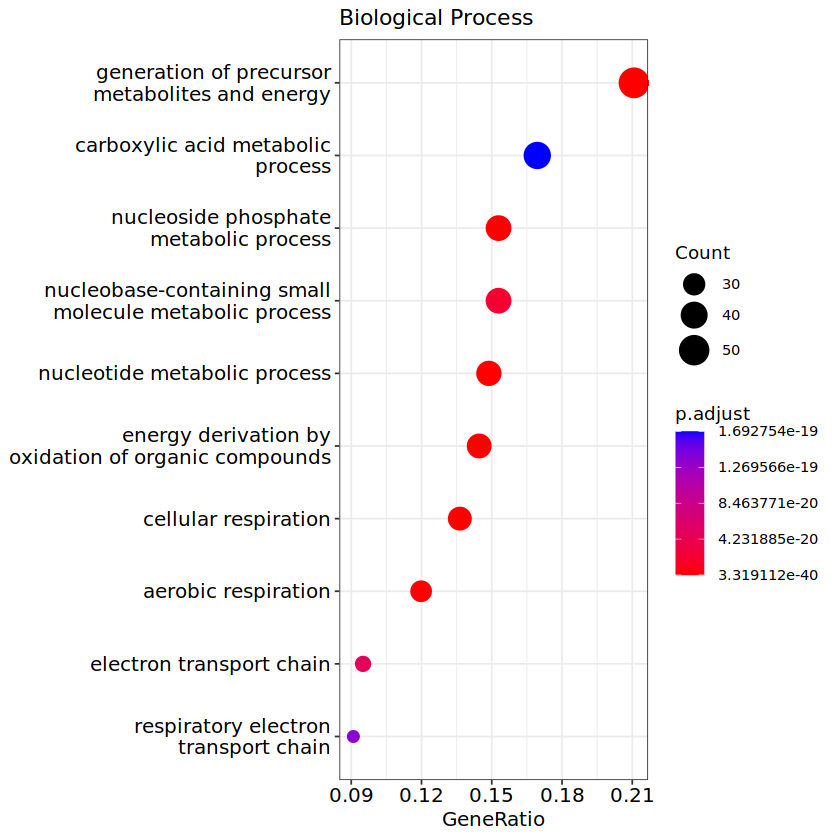

### Supplementary Material F4.png

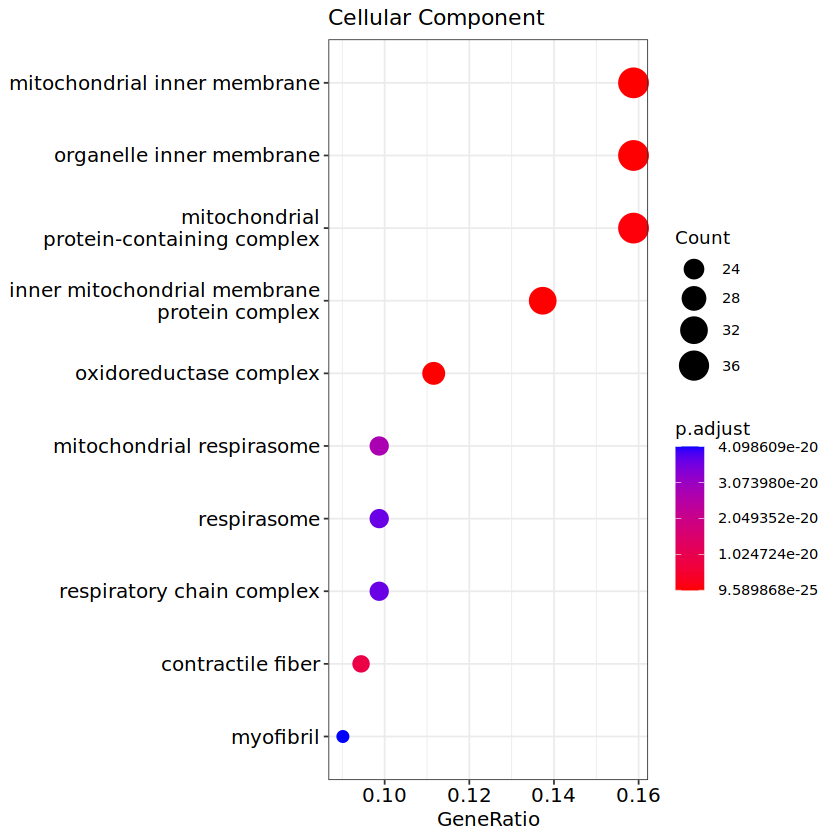

### Supplementary Material F5.png

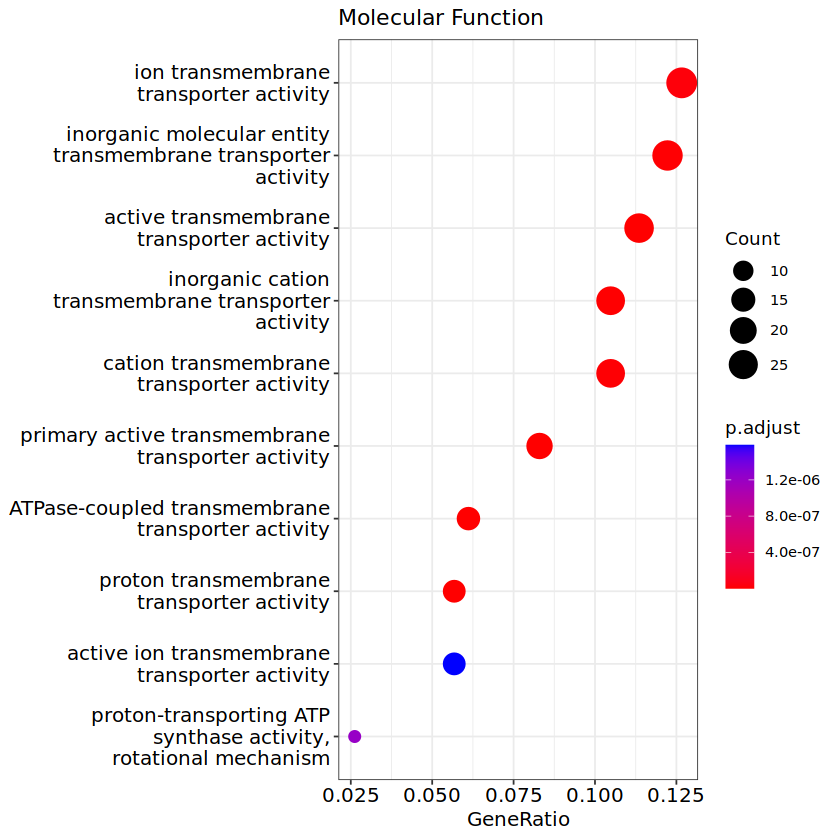
